# MicroRNA-378a-3p Modulates Inflammatory Responses of Keratinocytes to Atopic Dermatitis-Related Cytokines or *Staphylococcus aureus*

**DOI:** 10.64898/2026.03.16.711984

**Authors:** Kapilraj Periyasamy, Kristiina Kingo, Raül Habela Paneque, Anu Remm, Martin Pook, Helen Vaher, Külli Kingo, Ana Rebane

## Abstract

miR-378a-3p has been reported to be upregulated in the lesional skin of patients with atopic dermatitis (AD); however, its function in AD remains unclear. Here, we demonstrate that miR-378a-3p expression is induced by IL-4 and live *Staphylococcus aureus* (*S. aureus*) in normal human epidermal keratinocytes (NHEKs) cultured in proliferative conditions or in a 3D epidermal culture model. Transcriptomic profiling and gene set enrichment analysis of miR-378a-3p-transfected NHEKs revealed positive enrichment of inflammatory response pathways alongside downregulation of genes associated with epidermal development. More specifically, miR-378a-3p enhanced expression of multiple NF-κB-dependent inflammatory mediators, accompanied by increased phosphorylation of p65, indicating activation of canonical NF-κB pathway. Notably, miR-378a-3p concomitantly reduced the expression of several NF-κB family members and upstream adaptor molecules, supporting a model in which miR-378a-3p promotes canonical NF-κB activity through coordinated modulation of multiple components within the NF-κB regulatory network. In NHEKs exposed to live *S. aureus*, miR-378a-3p significantly increased the secretion of IL-1β, IL-1Ra, and IL-8, indicating that miR-378a-3p may amplify innate immune responses triggered by *S. aureus* colonization in AD. Collectively, these findings identify miR-378a-3p as a positive regulator of keratinocyte inflammatory responses that may contribute to AD exacerbation, particularly in the context of *S. aureus* colonization.

## Introduction

Keratinocytes are the major cell type in the epidermis, forming the physical barrier of the skin and actively responding to injury or pathogen invasion by producing cytokines, chemokines, and antimicrobial peptides. Dysregulated keratinocyte responses and impairment of the epithelial barrier contribute to the development of various skin diseases, including atopic dermatitis (AD) (Jiang et al., 2020; Simmons and Gallo, 2024). AD is a common, heterogeneous, chronic inflammatory skin disease associated with predisposing genetic factors and increased T helper 2 (Th2) immune responses (Tsoi et al., 2019; Schuler et al., 2024; Fyhrquist et al., 2025; Ye and Lai, 2025). Although Th2 cytokines predominate, AD-related skin inflammation can also involve activation of Nuclear Factor κB (NF-κB), IL-17/IL-22, IL-1 and Interferon-γ (IFN-γ) pathways, depending on disease endotype (Fyhrquist et al., 2025). Another hallmark of AD is increased colonization by *Staphylococcus aureus* (*S. aureus*), whose abundance on the skin has been shown to correlate with disease severity and progression (Lefèvre-Utile et al., 2022; Chehadeh et al., 2026). We have previously shown that *S. aureus* activates the inflammasome in keratinocytes, leading to increased secretion of IL-18 and IL-1β, which subsequently activate the NF-κB pathway, providing a potential mechanism linking *S. aureus* colonization to disease severity in AD (Vaher et al., 2023).

MicroRNAs (miRNAs) are short non-coding RNAs that regulate gene expression by binding to target mRNAs, thereby directing their degradation or translational repression (Jonas and Izaurralde, 2015; Wilczynska and Bushell, 2015). Several miRNAs, including miR-146a, miR-155, and miR-10a, have been implicated in AD pathogenesis by modulating T cell and keratinocyte responses (Sonkoly et al., 2010; Rebane et al., 2014; Vaher et al., 2019; Weidner et al., 2021). miR-378a-3p, encoded within the first intron of the *PPARGC1B* (Peroxisome Proliferator-Activated Receptor Gamma Coactivator 1 Beta) gene, has been studied primarily in metabolic and oncogenic contexts (Krist et al., 2015; Machado et al., 2020; Niu et al., 2020; Castellani et al., 2022; Qin et al., 2022). In the skin, miR-378a-3p has been shown to be upregulated in psoriasis, a chronic inflammatory skin disease characterized by IL-23/Th17 pathway activation and enhanced innate immune response (Griffiths et al., 2021; Soonthornchai et al., 2021; Xia et al., 2022). In keratinocytes, miR-378a-3p has been shown to be induced by IL-17A and to modulate the NF-κB pathway by targeting the *NFKBIA* mRNA, which encodes the NF-κB inhibitor IκBα (Xia et al., 2022). We have previously reported increased miR-378a-3p expression in lesional skin of both AD and psoriasis (Carreras-Badosa et al., 2022). However, the function of miR-378a-3p in AD, including in disease-related context in keratinocytes, remains to be studied.

In this study, we investigated the expression and function of miR-378a-3p in normal human primary keratinocytes (NHEKs) stimulated with AD-related pro-inflammatory cytokines and *S. aureus* infection. Our findings indicate that beyond its role in psoriasis, miR-378a-3p is involved in the regulation of inflammatory responses in AD.

## Results

### miR-378a-3p is induced by IL-4 and *S. aureus*, and modulates the transcriptional landscape of keratinocytes

To identify AD-related factors that regulate miR-378a-3p, we measured its expression in proliferating NHEKs grown in two-dimensional (2D) monolayers or three-dimensional (3D) epidermal equivalents. Consistent with previous findings (Xia et al., 2022), IL-17A induced miR-378a-3p expression (Figure 1a). Notably, IL-4 and live *Staphylococcus aureus* (*S. aureus*) also upregulated miR-378a-3p in both 2D and 3D NHEK cultures (Figure 1a). In addition, we observed slight downregulation of miR-378a-3p in the presence of IFN-γ. To investigate the function of miR-378a-3p in the context of AD, we transfected NHEKs with miR-378a-3p or negative control mimic followed by stimulation with IL-4, IL-17A, or IFN-γ and transcriptome and pathway analyses (Figure 1b, c). First, principal component analysis (PCA) was performed, which showed that cytokine stimulation produced essential transcriptomic shifts, with IFN-γ inducing the strongest response (Figure 1d), while overexpression of miR-378a-3p caused modest but distinguishable changes in each condition compared to negative control (Figure 1d, Supplementary Figure S1a-c). Next, we identified differentially expressed genes (DEGs; baseMean ≥ 15, FDR < 0.05) by comparing miR-378a-3p-transfected groups with corresponding control groups (Supplementary Table S1). In unstimulated cells, miR-378a-3p overexpression resulted in 3165 DEGs (1386 up, 1779 down). Cytokine stimulation further increased the number of miR-378a-3p-dependent DEGs, with the exception of IL-17A stimulation (Figure 1e). As shown in the UpSet plot, 334 genes were consistently upregulated and 538 downregulated across all four conditions regardless of cytokine stimulation (Figure 1f). The miR-378a-3p overexpression yielded the highest number of unique DEGs in cells stimulated with IFN-γ (1627 up, 1503 down), followed by IL-4 (664 up, 578 down), unstimulated condition (287 up, 285 down), and IL-17A (227 up, 178 down) (Figure 1f). Together, these data demonstrate that miR-378a-3p expression is modulated by different inflammatory cytokines, upregulated by *S. aureus* and exerts shared and cytokine-specific regulatory effects on the keratinocyte transcriptome, with the most pronounced impact observed in the presence of IFN-γ.

**Figure 1.**
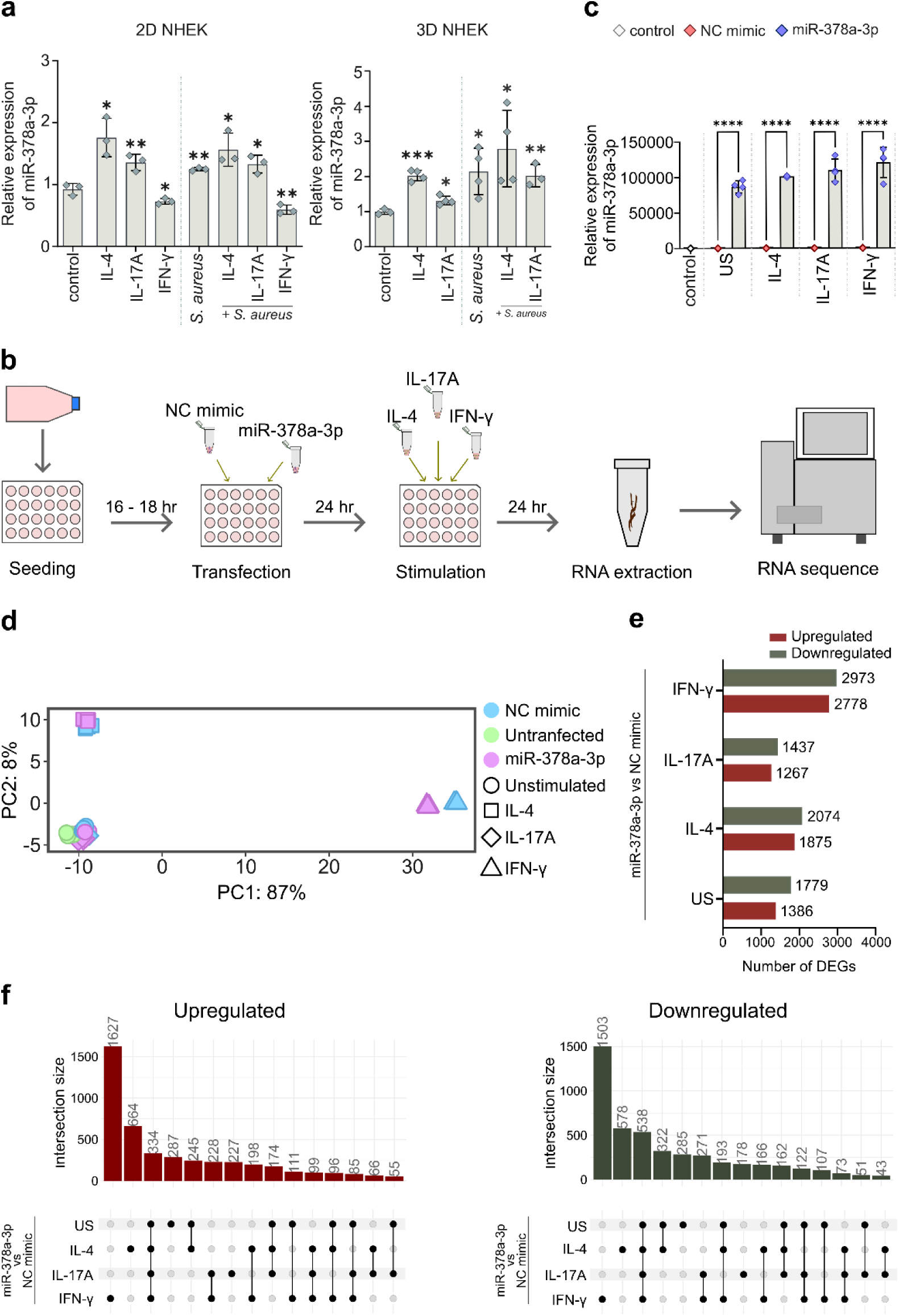
The expression regulation and influence of miR-378a-3p on the transcriptional landscape in NHEKs. (a, c) Relative expression of miR-378a-3p (RT-qPCR) in proliferating (2D) and 3D NHEK cultures stimulated with the indicated cytokines for 24 h or exposed to live *S. aureus* for 2 h, followed by medium change and sample collection 6 h later (a), or transfected with miR-378a-3p or negative control (NC) mimic in stimulated and unstimulated (US) conditions (c) as shown in (b). Control indicates untransfected and unstimulated conditions. Data are represented as mean ± SD (n = 3-4). Unpaired two-tailed Student’s *t*-test, \**p* < 0.05, \*\**p* < 0.01, \*\*\**p* < 0.001, \*\*\*\**p* < 0.0001. (b) Schematic overview of the RNA-seq workflow. (d) PCA of transcriptomic profiles. (e, f) The number of miR-378a-3p-influenced DEGs (baseMean ≥ 15, FDR < 0.05) (e) and UpSet plot showing unique and shared DEGs (f). Black dots represent unique DEGs in given condition, while black dots connected by lines represent DEGs shared between conditions.

### Overexpression of miR-378a-3p leads to gene expression changes linked with inflammatory pathways and epidermal development

To delineate gene networks influenced by miR-378a-3p, we next performed Gene Set Enrichment Analysis (GSEA) using both Hallmark and Gene Ontology (GO) Biological Process gene set collections. Analysis of Hallmark gene sets revealed positive enrichment of inflammation-related pathways, including inflammatory response, KRAS signaling, and TNF-α signaling via NF-κB, across all conditions, while p53 pathway and PI3K-AKT-mTOR signaling were enriched among downregulated genes (Figure 2a). Analysis of GO Biological Process datasets further showed positive enrichment of genes related to immune cell migration and chemokine response, whereas genes associated with skin epidermal development and differentiation were enriched among downregulated genes (Figure 2b). Despite influence on specific pathways, miR-378a-3p impact on global transcriptional landscape of cytokine responses was moderate (Supplementary Figure S2a). We next performed focused analyses of particular gene sets influenced by miR-378a-3p, while also considering the presence of its predicted targets among the DEGs selected by TargetScan (McGeary et al., 2019) (Supplementary Table S2). We first assessed the keratinocyte-specific NF-κB target gene set (MSigDB: HINATA_NFKB_TARGET_KERATINOCYTE_UP) and observed that several NF-κB-responsive genes were consistently upregulated across multiple conditions by miR-378a-3p, including *CXCL8*, *CCL20*, and *CXCL11*, as well as *ICAM1* (Figure 2c, d, extended list in Supplementary Figure S2b, c). Interestingly, among the downregulated NF-κB-dependent genes, IL-1 receptor antagonist gene, *IL1RN* was detected (Supplementary Figure S2b, c). When we examined other IL-1 family genes (Garlanda et al., 2013), we found that miR-378a-3p suppressed the expression of several IL-1 family cytokines (*IL1B*, *IL18*, *IL33*, *IL36G*) and receptor antagonists (*IL1RN* and *IL36RN*), while it upregulated the inhibitory receptor *SIGIRR* (Figure 2e, f). Among IL-1 family genes, *IL33* was detected as a predicted direct target (Supplementary Table S2), which has been experimentally validated before (Dubois-Camacho et al., 2019). Interestingly, even though we worked on proliferating keratinocytes, we observed negative enrichment of epidermal development terms (Figure 2b), and therefore carried out focused analysis for a relevant GO gene set (MSigDB: GOBP_EPIDERMIS_DEVELOPMENT). As shown in Supplementary Figure S3a, a large number of genes crucial for epidermal development were consistently downregulated by miR-378a-3p, including keratins (*KRT14, KRT15, KRT16*), desmosomal components (*PKP1, DSP, PPL*), and differentiation regulators (*TGM1, ST14*), while some of the differentiation markers were upregulated (*KRT10*). Among downregulated genes, *TGM1* was identified as a predicted target for miR-378a-3p (Supplementary Table S2). We also observed that miR-378a-3p overexpression reduced MHC class I gene expression across multiple conditions and increased MHC class II-related gene expression upon IFN-γ stimulation (Supplementary Figure S3b-d), consistent with the known ability of IFN-γ to induce MHC class II expression in keratinocytes (Albanesi et al., 1998). Collectively, transcriptomic analyses indicate that overexpression of miR-378a-3p modulates inflammatory pathways across multiple stimulation conditions in keratinocytes. Intriguingly, miR-378a-3p also influenced genes associated with epidermal development, generating the hypothesis that miR-378a-3p may modulate epidermal development or keratinocyte differentiation.

**Figure 2.**
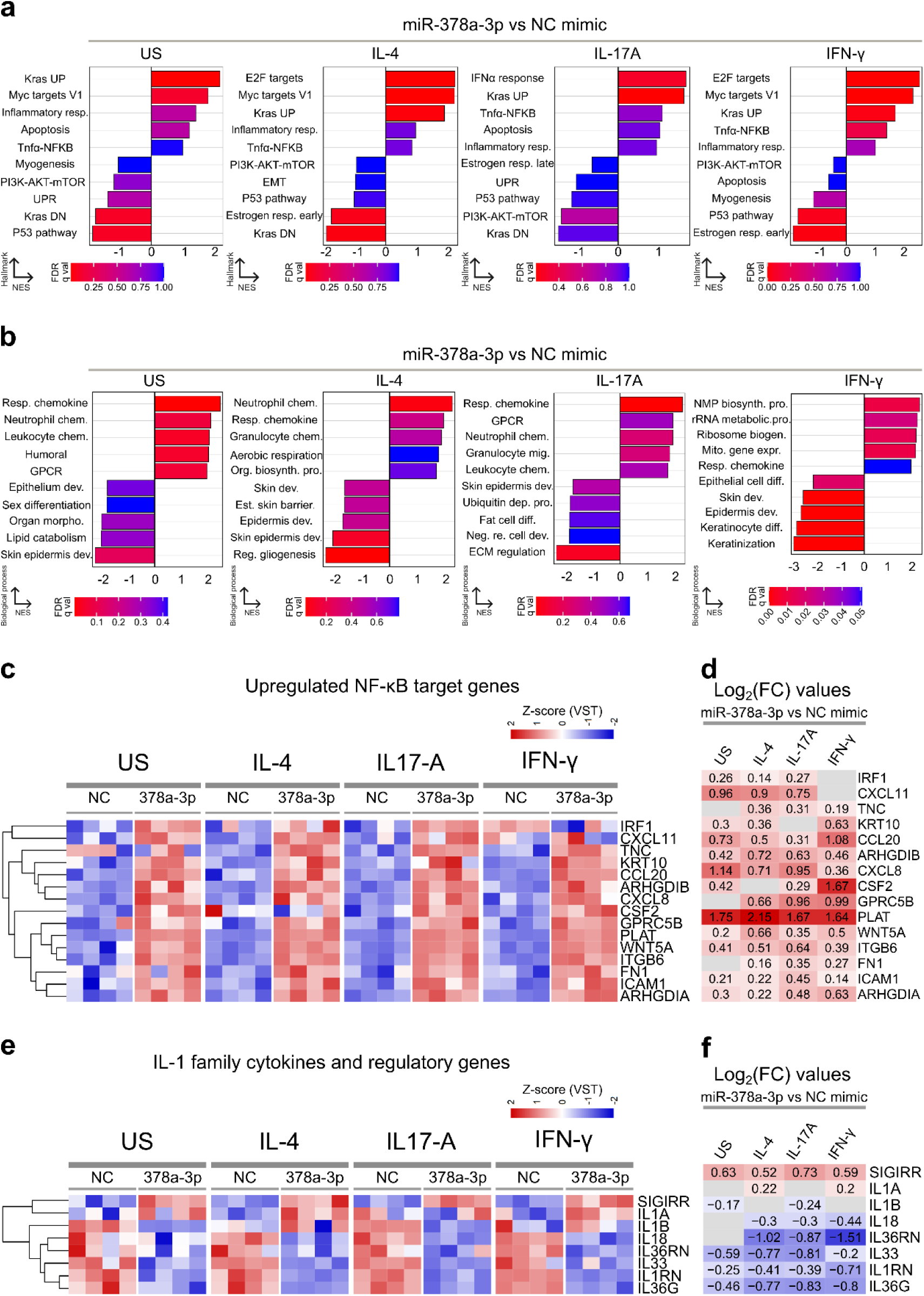
Transcriptomic signatures associated with miR-378a-3p overexpression in keratinocytes. (a, b) GSEA was performed on miR-378a-3p-regulated DEGs detected in unstimulated (US) and cytokine-stimulated conditions. Selected influenced Hallmark pathways (a) and GO biological processes (b) are shown. NES, normalized enrichment score. Full gene set names are provided in Supplementary Table S3. (c) Heatmap showing the expression changes (z-score) of consistently upregulated NF-κB-responsive genes from MSigDB. (d) Log_2_ fold-change values for the genes shown in (c). (e) Heatmap showing expression changes (z-score) of modulated IL-1 family members. (f) Log_2_ fold-change values for genes shown in (e). (d, f) Grey boxes indicate no change.

### Validation of miR-378a-3p-mediated regulation of inflammatory mediators

To validate miR-378a-3p effects detected by transcriptomic analysis, we measured the expression of selected inflammatory mediators at the mRNA level by RT-qPCR and at the protein level by ELISA. Consistent with the RNA-seq data (Figure 2c), miR-378a-3p overexpression significantly increased mRNA expression of NF-κB-regulated inflammatory mediators, including *CXCL8*, *CCL20, ICAM1* and *CXCL5* while the mRNA expression of IL-1 family genes, including *IL1B*, *IL18, IL1RN* and *IL36G* was downregulated in most of the conditions (Figure 3a, b and Supplementary Figure S4a). In line with this, miR-378a-3p overexpression significantly increased secretion of IL-8 (*CXCL8*) in most of the conditions, with the strongest effect observed following IL-17A stimulation (Figure 3c), while it decreased IL-1β and IL-1 receptor antagonist IL-1Ra protein levels in supernatants of NHEKs in all conditions (Figure 3d). We also observed that transfection of miR-378a-3p increased the proportion of HLA-DR-positive (MHC class II positive) cells from 26.9% to 42.7% following IFN-γ stimulation (Supplementary Figure S4b). Together, the validation experiments confirm that miR-378a-3p modulates the expression of inflammatory mediators in human primary keratinocytes, upregulating some while suppressing others.

**Figure 3.**
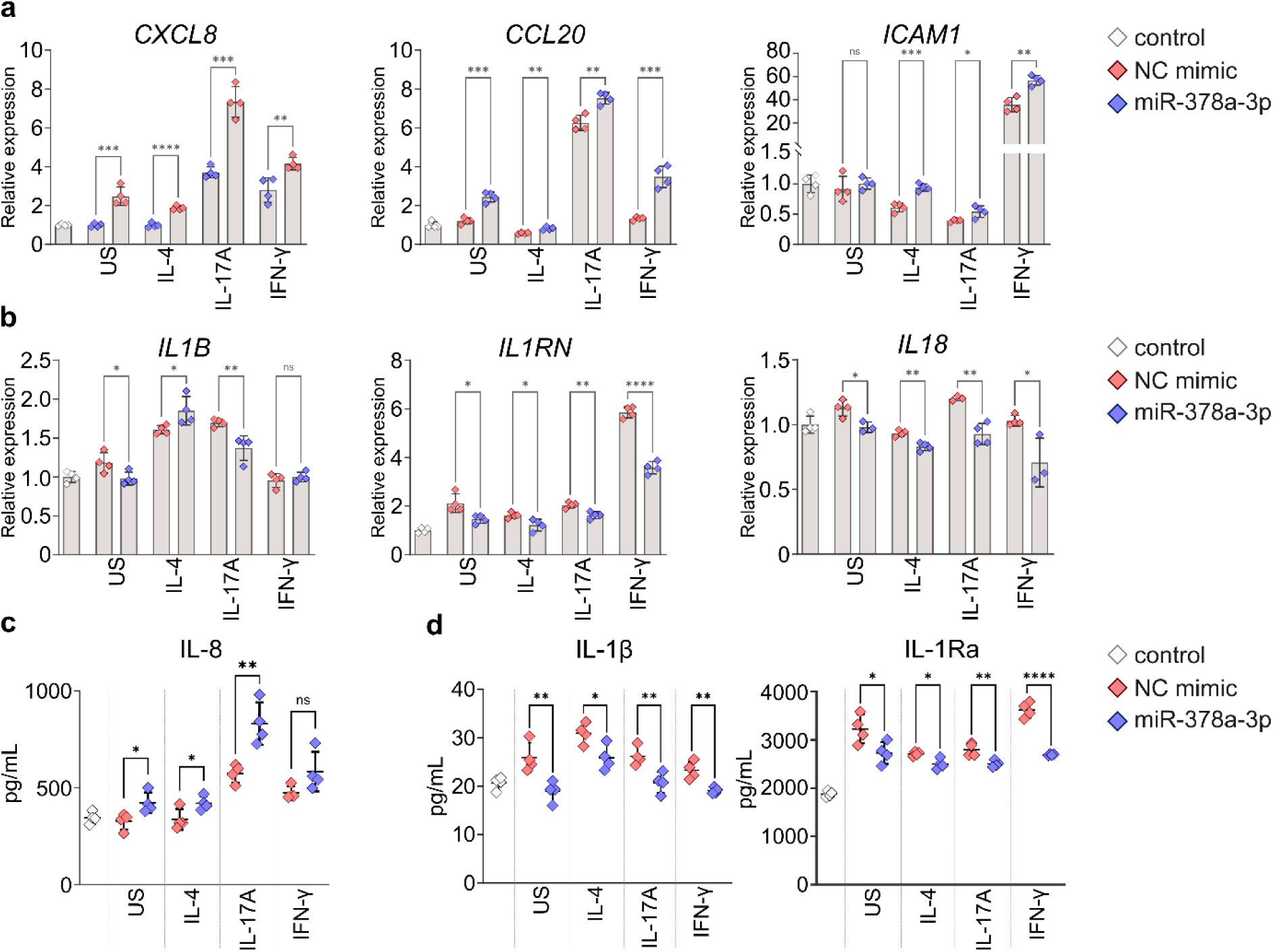
miR-378a-3p overexpression alters the expression of mediators of inflammatory response at mRNA and protein level. NHEKs were transfected with miR-378a-3p or negative control (NC) mimic for 24 h and then stimulated with the indicated cytokines for an additional 24 h. Untransfected and unstimulated NHEKs are designated as control. (a, b) RT-qPCR analysis of indicated genes. (c, d) Levels of IL-8, IL-1β, and IL-1Ra in cell culture supernatants measured by ELISA. (a-d) Data are presented as mean ± SD (n = 3-4). Unpaired two-tailed Student’s *t*-test. US, unstimulated; ns, not significant; \**p* < 0.05, \*\**p* < 0.01, \*\*\**p* < 0.001, \*\*\*\**p* < 0.0001.

### miR-378a-3p overexpression leads to increased phosphorylation of p65

As transcriptomic and validation experiments indicated that miR-378a-3p modulates the activation of the NF-κB pathway. To further examine miR-378a-3p effect on the NF-κB signaling network, we analyzed the expression of genes known to regulate the NF-κB signaling pathway (KEGG hsa04064). Across unstimulated and cytokine-stimulated conditions, miR-378a-3p overexpression altered the expression of multiple upstream regulators, including adaptors, kinases, inhibitors and NF-κB family transcription factors (Hayden and Ghosh, 2012) (Figure 4a, b). Among the transcripts downregulated by miR-378a-3p in at least one condition, multiple predicted miR-378a-3p direct targets were found, including *TRAF3*, *TRAF6*, *TOLLIP*, and NF-κB subunits *RELA* (p65) and *REL* (c-Rel) (Figure 4a, b and Supplementary Table S2). We also observed reduced expression of other genes not identified as predicted targets, such as *PRKCA*, *TRADD*, *IRAK4*, *PRKCZ* and *NFKBIZ* (IκBζ), atypical IκB family protein that modulates NF-κB (Hayden and Ghosh, 2012; Müller et al., 2018). Concurrently, miR-378a-3p upregulated *BCL10* and *CARD11*, components of the CARD11-BCL10-MALT1 (CBM) complex (Turvey et al., 2014) (Figure 4a, b). Since canonical NF-κB pathway activation is mainly regulated at the protein level, we next performed western blot analyses, which revealed that miR-378a-3p overexpression was associated with a trend toward reduced levels of p65, RelB, and c-Rel across multiple conditions (Figure 4c, d). In unstimulated condition, moderately reduced levels were also observed for NF-κB1 (p105 and p50) and NF-κB2 (p100) (Figure 4c, d). Notably, despite reduced abundance of the NF-κB subunits, the level of phosphorylated p65 (p-p65; Ser536), a hallmark of activated p65 (Yu et al., 2020) was increased, resulting in elevated p-p65 to total p65 ratio in all conditions (Figure 4d, e), suggesting activation of the canonical NF-κB pathway. Intriguingly, IκBα (*NFKBIA*), a previously validated target of miR-378a-3p and a major inhibitor of p65 (Xia et al., 2022), was not significantly affected by miR-378a-3p in our experiments (Figure 4a-d, Supplementary Table S1 and Supplementary Figure S4a). In summary, these findings indicate that miR-378a-3p reshapes the NF-κB regulatory network, which leads to the activation of the canonical pathway in keratinocytes.

**Figure 4.**
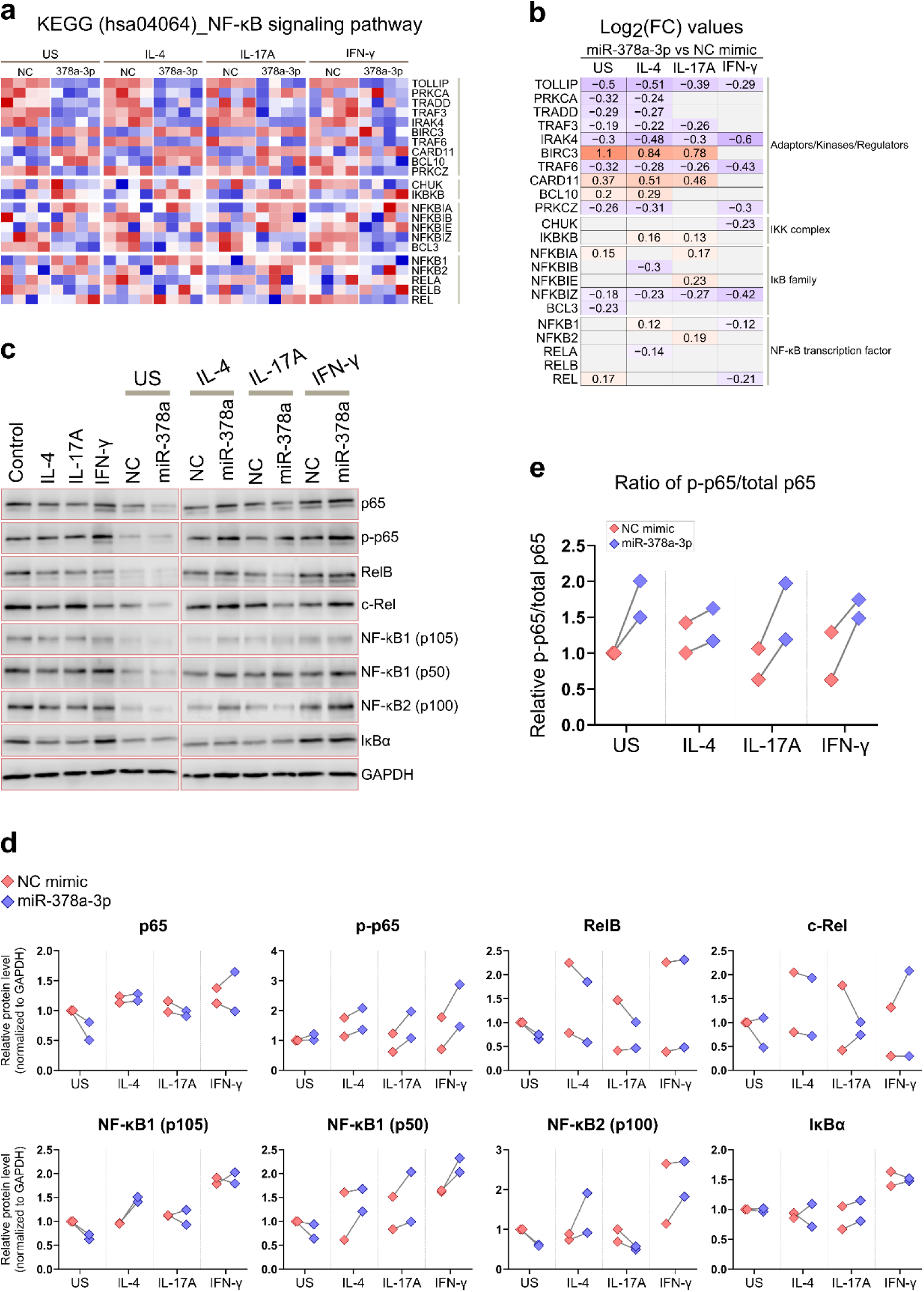
miR-378a-3p modulates the NF-κB pathway components at mRNA and protein levels. (a) Heatmap showing expression changes (z-score) of the NF-κB signaling pathway genes. (b) Log2 fold-change values for the genes shown in (a), grey boxes indicate no change. (c) Representative western blot images of key NF-κB pathway proteins. (d) Quantification of the NF-κB family protein levels. Normalized to GAPDH and presented relative to the unstimulated negative control (US NC, set to 1). (e) p-p65 and p65 band intensities were first normalized to GAPDH and the p-p65/total p65 ratio was then calculated relative to the US NC (set to 1). (d, e) Each symbol represents one independent experiment (n = 2), and are shown descriptively without statistical test. For each condition, values from NC and miR-378a-3p mimic within the same experiment are connected by lines.

### miR-378a-3p modulates *S. aureus*-induced inflammatory responses

*S. aureus* colonization is frequently associated with AD, contributing to disease exacerbation and severity (Park et al., 2013; Nakatsuji et al., 2016; Totté et al., 2016; Lefèvre-Utile et al., 2022). In keratinocytes, *S. aureus* activates the inflammasome, leading to the secretion of IL-1β, a strong upstream activator of canonical NF-κB pathway (Yu et al., 2020; Vaher et al., 2023). Therefore, we next investigated whether miR-378a-3p modulates keratinocyte responses to *S. aureus* infection. For that, we transfected NHEKs with miR-378a-3p or negative control mimic and subsequently infected cells with live *S. aureus* (Figure 5a). As expected, *S. aureus* infection alone strongly increased the mRNA expression of *CXCL8*, *CCL20,* and *IL1RN* (Figure 5b), and induced the secretion of IL-8, IL-1β, and IL-1Ra (Figure 5c). Notably, miR-378a-3p overexpression significantly amplified *S. aureus* response, leading to further increase in mRNA expression (*CXCL8*, *CCL20,* and *IL1RN*) and secretion of IL-1β, IL-1Ra and IL-8 compared to negative control mimic transfected condition (Figure 5b, c). These data indicate that miR-378a-3p exacerbates *S. aureus*-induced inflammatory responses in keratinocytes, suggesting that increased expression of miR-378a-3p in AD may intensify *S. aureus* induced innate immune activation.

**Figure 5.**
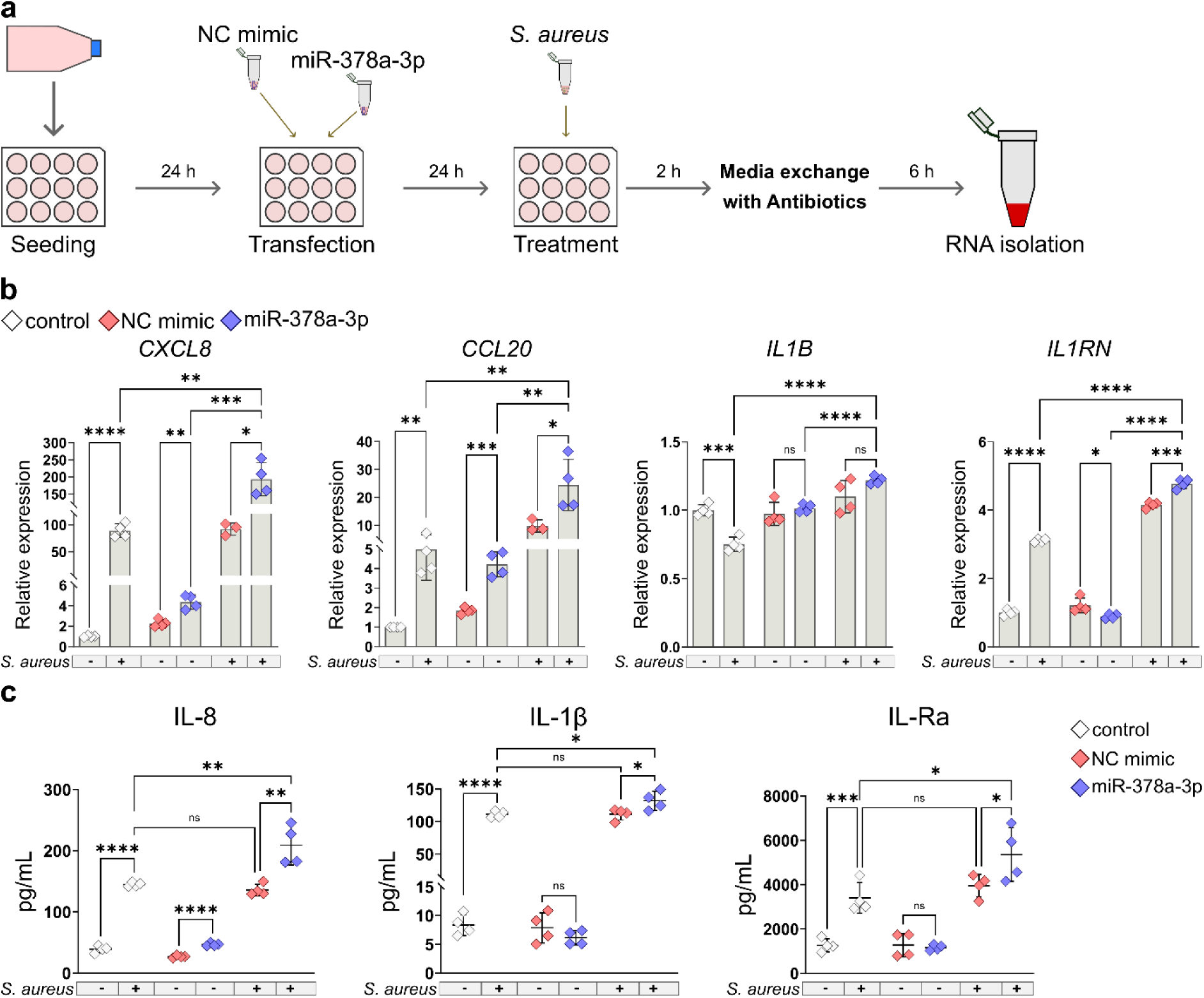
miR-378a-3p exacerbates *S. aureus*-induced inflammatory response in keratinocytes. (a) Schematic of the experimental workflow. NHEKs were transfected with miR-378a-3p or negative control (NC) for 24 h, followed by a 2 h infection with live *S. aureus* (1 × 10^7^ CFU/mL), and then additional 6 h incubation with media containing antibiotics. Control designates untransfected and untreated cells. (b) Relative mRNA expression of indicated inflammatory mediators, measured by RT-qPCR. (c) Concentration of cytokines in cell culture supernatants was measured by ELISA. (b, c) Data are presented as mean ± SD (n = 3-4). Unpaired two-tailed Student’s *t*-test. ns, not significant, \**p* < 0.05, \*\**p* < 0.01, \*\*\**p* < 0.001, \*\*\*\**p* < 0.0001.

## Discussion

MicroRNAs (miRNAs) are recognized as fine-tuning regulators of immune responses through their ability to modulate multiple or even a network of mRNAs, often within the same signaling pathway (Mehta and Baltimore, 2016; Weidner et al., 2021). In this study, we identify miR-378a-3p as a previously unrecognized regulator of keratinocyte responses relevant to AD. We show that miR-378a-3p is induced by IL-4 and live *S. aureus* in NHEKs and 3D epidermal models, linking type 2 inflammation and microbial exposure to miR-378a-3p regulation. Transcriptomic analysis and validation experiments further indicate that miR-378a-3p promotes canonical NF-κB-dependent inflammatory signaling, by reshaping the NF-κB regulatory network rather than acting through one dominant target mRNA. Notably, miR-378a-3p amplified cytokine release induced by *S. aureus*, suggesting a mechanism by which miR-378a-3p may amplify innate immune responses and skin inflammation in AD.

Previous studies have shown that miR-378a-3p is increased in the lesional skin of psoriasis patients and is induced by IL-17A stimulation in keratinocytes (Soonthornchai et al., 2021; Xia et al., 2022). Extending these findings, we show that miR-378a-3p is also induced by IL-4 and live *S. aureus* in NHEKs, which is consistent with our previous observation that miR-378a-3p is upregulated in the lesional skin of both AD and psoriasis patients (Carreras-Badosa et al., 2022).

Transcriptomic analysis revealed that miR-378a-3p overexpression had a relatively modest impact on global cytokine-induced gene expression changes; however, several pathways were consistently affected. These included among others positive enrichment of TNF-α signaling via NF-κB, which effect we validated by showing increase in IL-8 protein secretion and upregulation of *CXCL8*, *CCL20* and *ICAM1* mRNA levels. Interestingly, despite enhanced expression of NF-κB-dependent mediators, transcriptomic analysis revealed that several upstream regulators of NF-κB signaling were downregulated by miR-378a-3p, while NF-κB family transcription factors were largely unchanged. Notably, several of these genes, including *TRAF3, TRAF6, TOLLIP, RELA* (p65) and REL (c-Rel) are predicted targets of miR-378a-3p. In line with transcriptomic findings, western blot analysis showed a trend toward reduced abundance of p65, RelB, c-Rel, NF-κB1 (p105) and NF-κB2 (p100) in some conditions, particularly in unstimulated keratinocytes. Nevertheless, phosphorylated p65 (p-p65) and the p-p65/total p65 ratio were consistently increased across all conditions, indicating enhanced activation of canonical NF-κB signaling. In addition, miR-378a-3p increased expression of the CBM complex components, including *BCL10* and *CARD11*, which are known to promote NF-κB activation (Blonska and Lin, 2011; Yu et al., 2020; Qi et al., 2022). Interestingly, although miR-378a-3p has been reported to activate NF-κB signaling through targeting *NFKBIA* (IκBα) (Xia et al., 2022), we did not observe reduced *NFKBIA* mRNA or IκBα protein levels in our experimental system. Activation of canonical and non-canonical NF-κB pathways, is governed by complex regulatory dynamics involving differential kinetics, multiple feedback loops, and coordinated regulation of numerous pathway components (Hayden and Ghosh, 2012; Yu et al., 2020; Guo et al., 2024). In line with this framework, our results support a model in which miR-378a-3p enhances dominant canonical NF-κB signaling by reshaping multiple regulatory components rather than acting through a single dominant node, ultimately resulting in increased p65 activation.

Notably, the effect of miR-378a-3p on inflammatory responses was also evident when NHEKs were stimulated with live *S. aureus.* In this condition, miR-378a-3p enhanced *CXCL8*, *CCL20, IL1B* and *IL1RN* mRNA levels, and increased IL-8, IL-1β and IL-1Ra protein secretion. At the same time, in unstimulated conditions and in the presence of inflammatory cytokines alone, miR-378a-3p modestly reduced mRNA levels of IL-1 family genes and decreased secretion of IL-1β and IL-1Ra. Importantly, IL-1β secretion in these conditions was very low, indicating minimal or limited inflammasome activity. Notably, as increased IL-1β secretion is a hallmark of inflammasome activation and has previously been associated with increased disease severity in AD (Lefèvre-Utile et al., 2022; Vaher et al., 2023), our findings suggest that miR-378a-3p amplifies inflammatory responses in keratinocytes in AD particularly in case of *S. aureus* colonization.

Among other processes affected by miR-378a-3p, we observed that genes involved in epidermal differentiation and barrier integrity were downregulated, including *TGM1*, a predicted miR-378a-3p direct target and a key regulator in cornified envelope formation (Surbek et al., 2023; Ebrahimi Samani et al., 2024). Although it is interesting that this effect revealed in proliferating keratinocytes, to delineate whether miR-378a-3p contributes to barrier dysfunction through direct regulation of *TGM1*, further studies would be needed in 3D epidermal equivalents. Another interesting observation was that miR-378a-3p enhanced IFN-γ-induced MHC class II gene expression and increased the proportion of HLA-DR-positive keratinocytes. It has been suggested that keratinocytes expressing MHC class II are involved in the development of organized cellular clusters within the skin epithelium, which helps to recruit Th1 cells thereby maintaining cutaneous immune homeostasis and defense against pathogens (Tamoutounour et al., 2019). Whether this process is linked to skin inflammation in AD, remains to be studied.

Our study has also several limitations. First, miR-378a-3p function was assessed in proliferating keratinocytes, which precluded direct evaluation of its effect on epidermal differentiation and barrier formation. Second, although our data indicate that increased miR-378a-3p levels in AD are associated with enhanced activation of the canonical NF-κB signaling through modulation of multiple pathway components, the contribution of individual target genes was not resolved. While transcriptomic analyses support a multi-target mode of action, experimental identification of relevant direct miR-378a-3p targets would require additional large-scale approaches, such as Argonaute-CLIP-based methods, capturing miRNA–mRNA binding (Jaskiewicz et al., 2012), which are not optimized for NHEKs. In conclusion, our findings support a model in which increased miR-378a-3p, induced by both cytokine and microbial triggers, contributes to the modulation of inflammatory responses in AD. Further mechanistic studies will be required to delineate network of miR-378a-3p direct targets and to assess the functional relevance of in immune cells and in *in vivo* models of AD.

## Materials and Methods

### Cell culture experiments

Normal human epidermal keratinocytes (NHEKs; PromoCell, Heidelberg, Germany) were cultured in Keratinocyte Growth Medium 3 (KGM3; PromoCell) supplemented according to the manufacturer’s instructions. NHEKs were transfected with 60 nM miR-378a-3p mimic (Horizon Discovery, Cambridge, UK) or negative control (NC) mimic (Horizon Discovery) using cell-penetrating peptide PepFect14 (PF14) for 24 hours, as previously described (Vaher et al., 2023). Stimulation with IL-4 (40 ng/mL), IL-17A (10 ng/mL), or IFN-γ (20 ng/mL) (PeproTech, Rocky Hill, NJ, USA) was performed for 24 hours. Infection with *S. aureus* (strain DSM 2569) was done at 1 × 10^7^ CFU/mL, as previously described (Vaher et al., 2023). Detailed protocols are provided in the Supplementary Material and Methods.

### RNA extraction and RT-qPCR

Total RNA was isolated using the miRNeasy mini kit (Qiagen, Venlo, Netherlands) according to the manufacturer’s instructions. For mRNA analysis, cDNA was synthesized from 400–600 ng total RNA using the oligo-dT (TAG Copenhagen, Frederiksberg, Denmark) and qPCR was performed using HOT FIREPol EvaGreen qPCR Supermix (Solis Biodyne, Tartu, Estonia). Primer sequences are listed in Supplementary Table S4. miRNA expression was analyzed using the TaqMan MicroRNA Reverse Transcription Kit (Thermo Fisher Scientific) and TaqMan MicroRNA Assays. Detailed protocols are provided in the Supplementary Material and Methods.

### RNA sequencing, pathway and target analysis

RNA sequencing was performed on Illumina NovaSeq X Plus platform (PE150; 6 G raw data per sample) by Novogene (Cambridge, UK). Subsequent data analysis was performed on the Galaxy platform. Differentially expressed genes (DEGs) were defined as those with a FDR < 0.05 and DESeq2 baseMean ≥ 15. Gene Set Enrichment Analysis (GSEA v4.3.3) was conducted in pre-ranked mode. Predicted miR-378a-3p target genes were obtained from TargetScan (release 8.0) and are listed in Supplementary Table S2. Detailed protocols are provided in the Supplementary Material and Methods.

### Protein expression analyses

Secreted IL-8 (BioLegend, San Diego, CA, USA), IL-1β (BioLegend), and IL-1Ra (R&D Systems, Minneapolis, MN, USA) in culture supernatants were quantified using ELISA according to the manufacturers’ instructions. For western blotting, cells were lysed in RIPA buffer supplemented with cOmplete^TM^ protease inhibitor cocktail (Roche, Basel, Switzerland). The following primary antibodies were used: p65 (clone D14E12), phosphorylated p65 (p-p65; clone 93H1), RelB (clone D7D7W), c-Rel (clone E8Z5Y), NF-κB1 (p105/p50; clone 5D10D11), NF-κB2 (p100/p52; clone 18D10), IκBα (clone L35A5) (Cell Signaling Technology, Danvers, MA, USA), and GAPDH (clone 6C5; Santa Cruz Biotechnology, Dallas, TX, USA). For flow cytometry, cells were incubated with Fc receptor blocking reagent (Miltenyi Biotec, Bergisch Gladbach, Germany) and stained with FITC-conjugated anti-human HLA-DR (BioLegend; clone L243). Detailed protocols are provided in the Supplementary Material and Methods.

### Statistical analysis and visualization

Statistical analysis of RT-qPCR and ELISA results was performed using Prism 10 (GraphPad Software). Statistical significance was determined using an unpaired two-tailed Student’s t-test considering that data from independent conditions were compared, p-value ≤ 0.05 was considered statistically significant. Data are presented as mean ± SD. Volcano plots, heatmaps, sample-to-sample distance matrices, and log2 fold-change visualizations were generated using R (RStudio 2025.05.1+513). Detailed protocols are provided in the Supplementary Material and Methods.

## Data Availability Statement

Additional data are available from the corresponding author upon request.

## Acknowledgements

This research was supported by the Estonian Research Council grants PRG1259 and PRG1189.

## Author contributions

Conceptualization: KP, MP, HV, AReb

Data Curation: KP, KrK, RHP, ARem

Formal Analysis: KP, KrK, RHP, ARem

Investigation: KP, KrK, RHP, ARem, MP

Methodology: KP, RHP, MP, AReb

Supervision and funding acquisition: KüK, AReb

Writing – Original Draft preparation: KP, AReb

Writing – Review and Editing: KP, RHP, MP, KüK, AReb

## Abbreviations

AD: atopic dermatitis
miR-378a-3p: microRNA-378a-3p
*S. aureus*: *Staphylococcus aureus*
NHEKs: normal human epidermal keratinocytes
NF-κB: nuclear factor κB
MHC: major histocompatibility complex
GSEA: gene set enrichment analysis

## References

Albanesi C, Cavani A, Girolomoni G. Interferon-γ-Stimulated Human Keratinocytes Express the Genes Necessary for the Production of Peptide-Loaded MHC Class II Molecules. J Invest Dermatol 1998;110:138–42.

Blonska M, Lin X. NF-κB signaling pathways regulated by CARMA family of scaffold proteins. Cell Res 2011;21:55–70.

Carreras-Badosa G, Maslovskaja J, Vaher H, Pajusaar L, Annilo T, Lättekivi F, et al. miRNA expression profiles of the perilesional skin of atopic dermatitis and psoriasis patients are highly similar. Sci Rep 2022;12:22645.

Castellani G, Buccarelli M, Lulli V, Ilari R, De Luca G, Pedini F, et al. MiR-378a-3p Acts as a Tumor Suppressor in Colorectal Cancer Stem-Like Cells and Affects the Expression of MALAT1 and NEAT1 lncRNAs. Front Oncol 2022;12:867886.

Chehadeh C, Nakatsuji T, Gallo RL. Staphylococcus aureus in Atopic Dermatitis: How a common bacterium exploits and drives disease. J Allergy Clin Immunol 2026.

Dubois-Camacho K, Diaz-Jimenez D, De la Fuente M, Quera R, Simian D, Martínez M, et al. Inhibition of miR-378a-3p by Inflammation Enhances IL-33 Levels: A Novel Mechanism of Alarmin Modulation in Ulcerative Colitis. Front Immunol 2019;10:2449.

Ebrahimi Samani S, Tatsukawa H, Hitomi K, Kaartinen MT. Transglutaminase 1: Emerging Functions beyond Skin. Int J Mol Sci 2024;25:10306.

Fyhrquist N, Yang Y, Karisola P, Alenius H. Endotypes of atopic dermatitis. J Allergy Clin Immunol 2025;156:24–40.e4.

Garlanda C, Dinarello CA, Mantovani A. The interleukin-1 family: back to the future. Immunity 2013;39:1003–18.

Griffiths CEM, Armstrong AW, Gudjonsson JE, Barker JNWN. Psoriasis. Lancet 2021;397:1301–15.

Guo Q, Jin Y, Chen X, Ye X, Shen X, Lin M, et al. NF-κB in biology and targeted therapy: new insights and translational implications. Signal Transduct Target Ther 2024;9:53.

Hayden MS, Ghosh S. NF-κB, the first quarter-century: remarkable progress and outstanding questions. Genes Dev 2012;26:203–34.

Jaskiewicz L, Bilen B, Hausser J, Zavolan M. Argonaute CLIP--a method to identify in vivo targets of miRNAs. Methods 2012;58:106–12.

Jiang Y, Tsoi LC, Billi AC, Ward NL, Harms PW, Zeng C, et al. Cytokinocytes: the diverse contribution of keratinocytes to immune responses in skin. JCI Insight 2020;5:e142067.

Jonas S, Izaurralde E. Towards a molecular understanding of microRNA-mediated gene silencing. Nat Rev Genet 2015;16:421–33.

Krist B, Florczyk U, Pietraszek-Gremplewicz K, Józkowicz A, Dulak J. The Role of miR-378a in Metabolism, Angiogenesis, and Muscle Biology. Int J Endocrinol 2015;2015:281756.

Lefèvre-Utile A, Saichi M, Oláh P, Delord M, Homey B, Soumelis V. Transcriptome-based identification of novel endotypes in adult atopic dermatitis. Allergy 2022;77:1486–98.

Machado IF, Teodoro JS, Palmeira CM, Rolo AP. miR-378a: a new emerging microRNA in metabolism. Cell Mol Life Sci 2020;77:1947–58.

McGeary SE, Lin KS, Shi CY, Pham TM, Bisaria N, Kelley GM, et al. The biochemical basis of microRNA targeting efficacy. Science 2019;366:eaav1741.

Mehta A, Baltimore D. MicroRNAs as regulatory elements in immune system logic. Nat Rev Immunol 2016;16:279–94.

Müller A, Hennig A, Lorscheid S, Grondona P, Schulze-Osthoff K, Hailfinger S, et al. IκBζ is a key transcriptional regulator of IL-36-driven psoriasis-related gene expression in keratinocytes. Proc Natl Acad Sci U S A 2018;115:10088–93.

Nakatsuji T, Chen TH, Two AM, Chun KA, Narala S, Geha RS, et al. Staphylococcus aureus Exploits Epidermal Barrier Defects in Atopic Dermatitis to Trigger Cytokine Expression. J Invest Dermatol 2016;136:2192–200.

Niu F, Dzikiewicz-Krawczyk A, Koerts J, de Jong D, Wijenberg L, Fernandez Hernandez M, et al. MiR-378a-3p Is Critical for Burkitt Lymphoma Cell Growth. Cancers (Basel) 2020;12:3546.

Park H-Y, Kim C-R, Huh I-S, Jung M-Y, Seo E-Y, Park J-H, et al. Staphylococcus aureus Colonization in Acute and Chronic Skin Lesions of Patients with Atopic Dermatitis. Ann Dermatol 2013;25:410–6.

Qi T, Luo Y, Cui W, Zhou Y, Ma X, Wang D, et al. Crosstalk between the CBM complex/NF-κB and MAPK/P27 signaling pathways of regulatory T cells contributes to the tumor microenvironment. Front Cell Dev Biol 2022;10:911811.

Qin Y, Liang R, Lu P, Lai L, Zhu X. Depicting the Implication of miR-378a in Cancers. Technol Cancer Res Treat 2022;21:15330338221134384.

Rebane A, Runnel T, Aab A, Maslovskaja J, Rückert B, Zimmermann M, et al. MicroRNA-146a alleviates chronic skin inflammation in atopic dermatitis through suppression of innate immune responses in keratinocytes. J Allergy Clin Immunol 2014;134:836–847.e11.

Schuler CF 4th, Tsoi LC, Billi AC, Harms PW, Weidinger S, Gudjonsson JE. Genetic and Immunological Pathogenesis of Atopic Dermatitis. J Invest Dermatol 2024;144:954–68.

Simmons J, Gallo RL. The Central Roles of Keratinocytes in Coordinating Skin Immunity. J Invest Dermatol 2024;144:2377–98.

Sonkoly E, Janson P, Majuri M-L, Savinko T, Fyhrquist N, Eidsmo L, et al. MiR-155 is overexpressed in patients with atopic dermatitis and modulates T-cell proliferative responses by targeting cytotoxic T lymphocyte–associated antigen 4. J Allergy Clin Immunol 2010;126:581–589.e20.

Soonthornchai W, Tangtanatakul P, Meesilpavikkai K, Dalm V, Kueanjinda P, Wongpiyabovorn J. MicroRNA-378a-3p is overexpressed in psoriasis and modulates cell cycle arrest in keratinocytes via targeting BMP2 gene. Sci Rep 2021;11:14186.

Surbek M, Van de Steene T, Sachslehner AP, Golabi B, Griss J, Eyckerman S, et al. Cornification of keratinocytes is associated with differential changes in the catalytic activity and the immunoreactivity of transglutaminase-1. Sci Rep 2023;13:21550.

Tamoutounour S, Han S-J, Deckers J, Constantinides MG, Hurabielle C, Harrison OJ, et al. Keratinocyte-intrinsic MHCII expression controls microbiota-induced Th1 cell responses. Proc Natl Acad Sci U S A 2019;116:23643–52.

Totté JEE, van der Feltz WT, Hennekam M, van Belkum A, van Zuuren EJ, Pasmans SGMA. Prevalence and odds of Staphylococcus aureus carriage in atopic dermatitis: a systematic review and meta-analysis. Br J Dermatol 2016;175:687–95.

Tsoi LC, Rodriguez E, Degenhardt F, Baurecht H, Wehkamp U, Volks N, et al. Atopic Dermatitis Is an IL-13-Dominant Disease with Greater Molecular Heterogeneity Compared to Psoriasis. J Invest Dermatol 2019;139:1480–9.

Turvey SE, Durandy A, Fischer A, Fung S-Y, Geha RS, Gewies A, et al. The CARD11-BCL10-MALT1 (CBM) signalosome complex: Stepping into the limelight of human primary immunodeficiency. J Allergy Clin Immunol 2014;134:276–84.

Vaher H, Kingo Kristiina, Kolberg P, Pook M, Raam L, Laanesoo A, et al. Skin Colonization with S. aureus Can Lead to Increased NLRP1 Inflammasome Activation in Patients with Atopic Dermatitis. J Invest Dermatol 2023;143:1268–1278.e8.

Vaher H, Runnel T, Urgard E, Aab A, Carreras Badosa G, Maslovskaja J, et al. miR-10a-5p is increased in atopic dermatitis and has capacity to inhibit keratinocyte proliferation. Allergy 2019;74:2146–56.

Weidner J, Bartel S, Kılıç A, Zissler UM, Renz H, Schwarze J, et al. Spotlight on microRNAs in allergy and asthma. Allergy 2021;76:1661–78.

Wilczynska A, Bushell M. The complexity of miRNA-mediated repression. Cell Death Differ 2015;22:22–33.

Xia P, Pasquali L, Gao C, Srivastava A, Khera N, Freisenhausen JC, et al. miR-378a regulates keratinocyte responsiveness to interleukin-17A in psoriasis. Br J Dermatol 2022;187:211–22.

Ye J, Lai Y. Keratinocytes: new perspectives in inflammatory skin diseases. Trends Mol Med 2025;31:1103–13.

Yu H, Lin L, Zhang Z, Zhang H, Hu H. Targeting NF-κB pathway for the therapy of diseases: mechanism and clinical study. Signal Transduct Target Ther 2020;5:209.

