## Supplementary Information (Methods, Figures, Tables) for "MicroRNA-378a-3p Modulates Inflammatory Responses of Keratinocytes to Atopic Dermatitis-Related Cytokines or *Staphylococcus aureus*"

**Supplementary Material and Methods**

**Cell culture**

Normal human epidermal keratinocytes (NHEKs; PromoCell, Heidelberg, Germany) were cultured in Keratinocyte Growth Medium 3 (KGM3; PromoCell) supplemented according to the manufacturer’s instructions, with the addition of 5 U/mL of penicillin-streptomycin (Thermo Fisher Scientific, Waltham, MA, USA). For three-dimensional (3D) culture, 5 × 10^4^ NHEKs were seeded onto ThinCert cell culture inserts (Greiner Bio-One, Kremsmünster, Austria) and cultured at the air-liquid interface as previously described (Vaher et al., 2023), using a 1:1 mixture of KGM3 (PromoCell) and DMEM (Thermo Fisher Scientific).

**miRNA transfection and cytokine stimulation**

For transfection experiments, NHEKs were seeded at a density of 4 × 10^4^ cells/well in 24-well plates or 7 × 10^4^ cells/well in 12-well plates. After 16-18 hours (h), cells were transfected with 60 nM miR-378a-3p mimic (Horizon Discovery, Cambridge, UK) or negative control (NC) mimic (Horizon Discovery) using cell-penetrating peptide PepFect14 (PF14) as previously described (Vaher et al., 2023). Briefly, complexes of miRNA:PF14 at a 1:17 molar ratio (miRNA:PF14) were prepared in Milli-Q water at 1/10 of the final transfection volume, incubated for 1 h at room temperature, and then added to the culture medium. When indicated, after 24 h of incubation, cytokines were added for an additional 24 h before harvesting cells and/or supernatants. For experiments involving cytokine stimulation alone, cytokines were added to cells 16-18 h after seeding for 24 h. The following cytokine concentrations were used: IL-4 (40 ng/mL), IL-17A (10 ng/mL), and IFN-γ (20 ng/mL) (PeproTech, Rocky Hill, NJ, USA).

***S. aureus* infection**

Cells were infected with live *S. aureus* (strain DSM 2569), obtained from the Core Facility for Biosafety (University of Tartu, Tartu, Estonia), at a multiplicity of infection of approximately 100 (corresponding to 1 × 10^7^ CFU/mL) as previously described (Vaher et al., 2023). After 2 h, the medium was removed, cells were washed twice with PBS containing 50 U/mL penicillin-streptomycin, and fresh KGM3 containing 5 U/mL of penicillin-streptomycin was added for an additional 6 h. When indicated, NHEKs were transfected with negative control or miR-378a-3p mimic as described above and incubated for 24 h before infection.

**RNA extraction and RT-qPCR**

Total RNA was isolated using the miRNeasy Mini kit (Qiagen, Venlo, Netherlands) with on-column DNase treatment (RNase-Free DNase Set; Qiagen) according to the manufacturer’s instructions. For mRNA analysis, cDNA was synthesized from 400–600 ng total RNA using the oligo-dT (TAG Copenhagen, Frederiksberg, Denmark), dNTP mix, RevertAid reverse transcriptase, and RiboLock RNase inhibitor (Thermo Fisher Scientific). qPCR was performed using HOT FIREPol EvaGreen qPCR Supermix (Solis Biodyne, Tartu, Estonia) on a QuantStudio 12K Flex system (Thermo Fisher Scientific). Relative expression was calculated using the ΔΔCt method with *EEF1A1* as the reference gene. Primer sequences are listed in Supplementary Table S4. miRNA expression was analyzed using the TaqMan MicroRNA Reverse Transcription Kit (Thermo Fisher Scientific) and TaqMan MicroRNA Assays for miR-378a-3p (Thermo Fisher Scientific), normalized to let-7a (Thermo Fisher Scientific).

**RNA sequencing and pathway analysis**

RNA sequencing was performed using a directional mRNA library preparation protocol with poly(A) enrichment and sequenced on an Illumina NovaSeq X Plus platform (PE150; 6 G raw data per sample) (Novogene, Cambridge, UK). Subsequent data analysis was conducted on the Galaxy platform. Quality control was performed with FastQC (v0.11.8), reads were trimmed with Cutadapt (v4.4), and mapped with STAR (v2.7.8a) using the GRCh38 as the reference genome. Read counting was performed with featureCounts (v2.0.3) and differential expression analysis was conducted using DESeq2 (v2.11.40.6). Differentially expressed genes (DEGs) were defined as those with a FDR < 0.05 and DESeq2 baseMean ≥ 15.

Gene Set Enrichment Analysis (GSEA v4.3.3) was performed using the pre-ranked method. Genes were ranked by log_2_ fold change values to generate the ranked lists for GSEA. Annotated gene sets were obtained from the Molecular Signatures Database (MSigDB). Additionally, we used NF-κB pathway curated genes from KEGG pathway (hsa04064) (Figure 4a), and IL-1 family genes (Figure 2e) based on the literature (Garlanda et al., 2013).

Predicted miR-378a-3p target genes were obtained from TargetScan (release 8.0). Both targets with conserved and non-conserved sites were included. The predicted targets were intersected with DEGs (baseMean ≥ 15, FDR < 0.05, classified to upregulated or downregulated) from each condition (unstimulated, IL-4, IL-17A, and IFN-γ-stimulated). TargetScan-predicted targets that were significantly regulated in the same direction across all four conditions were identified and classified as shared regulated predicted targets (Supplementary Table S2).

**ELISA**

Levels of secreted IL-8 (BioLegend, San Diego, CA, USA), IL-1β (BioLegend), and IL-1Ra (R&D Systems, Minneapolis, MN, USA) in culture supernatants were quantified using commercial ELISA kits according to the manufacturers’ instructions. Absorbance was measured using a Ledetect 96 microplate reader (Labexim Products, Lengau, Austria), and concentrations were calculated using a four-parameter logistic (4PL) regression model (MyAssays.com).

**Western blotting**

Cells were lysed in RIPA buffer supplemented with cOmplete^TM^ protease inhibitor cocktail (Roche, Basel, Switzerland) according to the manufacturer’s instructions. Protein samples were resolved on 10% SDS-PAGE gels and transferred to PVDF membranes. Membranes were incubated overnight at 4 °C with primary antibodies against p65 (clone D14E12), phosphorylated p65 (p-p65; clone 93H1), RelB (clone D7D7W), c-Rel (clone E8Z5Y), NF-κB1 (p105/p50; clone 5D10D11), NF-κB2 (p100/p52; clone 18D10), IκBα (clone L35A5) (Cell Signaling Technology, Danvers, MA, USA), as well as GAPDH (clone 6C5; Santa Cruz Biotechnology, Dallas, TX, USA). HRP-conjugated secondary antibodies (goat anti-rabbit IgG or horse anti-mouse IgG; Cell Signaling Technology) were added for 1 h at room temperature. Images were quantified using Image Studio Software v6 (LI-COR Biosciences, Lincoln, NE, USA). The bands were selected with identical size across all samples, with median background subtraction (border width set to 3). Band densities were normalized to GAPDH, and expressed relative to negative control mimic, as indicated in the figure legends.

**Flow cytometry**

Cells were detached using 4 mM EDTA (15 min at 37 °C), washed, and resuspended in FACS buffer (PBS containing 0.5% BSA and 2 mM EDTA) and incubated with Fc receptor blocking reagent (Miltenyi Biotec, Bergisch Gladbach, Germany) for 10 min on ice. Subsequent antibody staining was performed using cold FACS buffer, and cells were kept on ice. Cells were stained with FITC-conjugated anti-human HLA-DR antibody (BioLegend; clone L243) for 30 min and washed twice with cold FACS buffer. Five minutes before acquisition, 0.5 µg/mL of DAPI was added to the samples. Data were acquired on a BD LSRFortessa cell analyzer (BD Biosciences, San Jose, CA, USA) and analyzed using Floreada.io software.

**Statistical analysis and visualization**

Volcano plots, heatmaps, sample-to-sample distance matrices, and log_2_ fold-change visualizations were generated using R (RStudio 2025.05.1+513). Statistical analysis was performed using Prism 10 (GraphPad Software). Data are presented as mean ± SD. Statistical significance was determined using an unpaired two-tailed Student’s *t*-test considering that data from independent conditions were compared. A *p*-value ≤ 0.05 was considered statistically significant.

**Supplementary Figures**

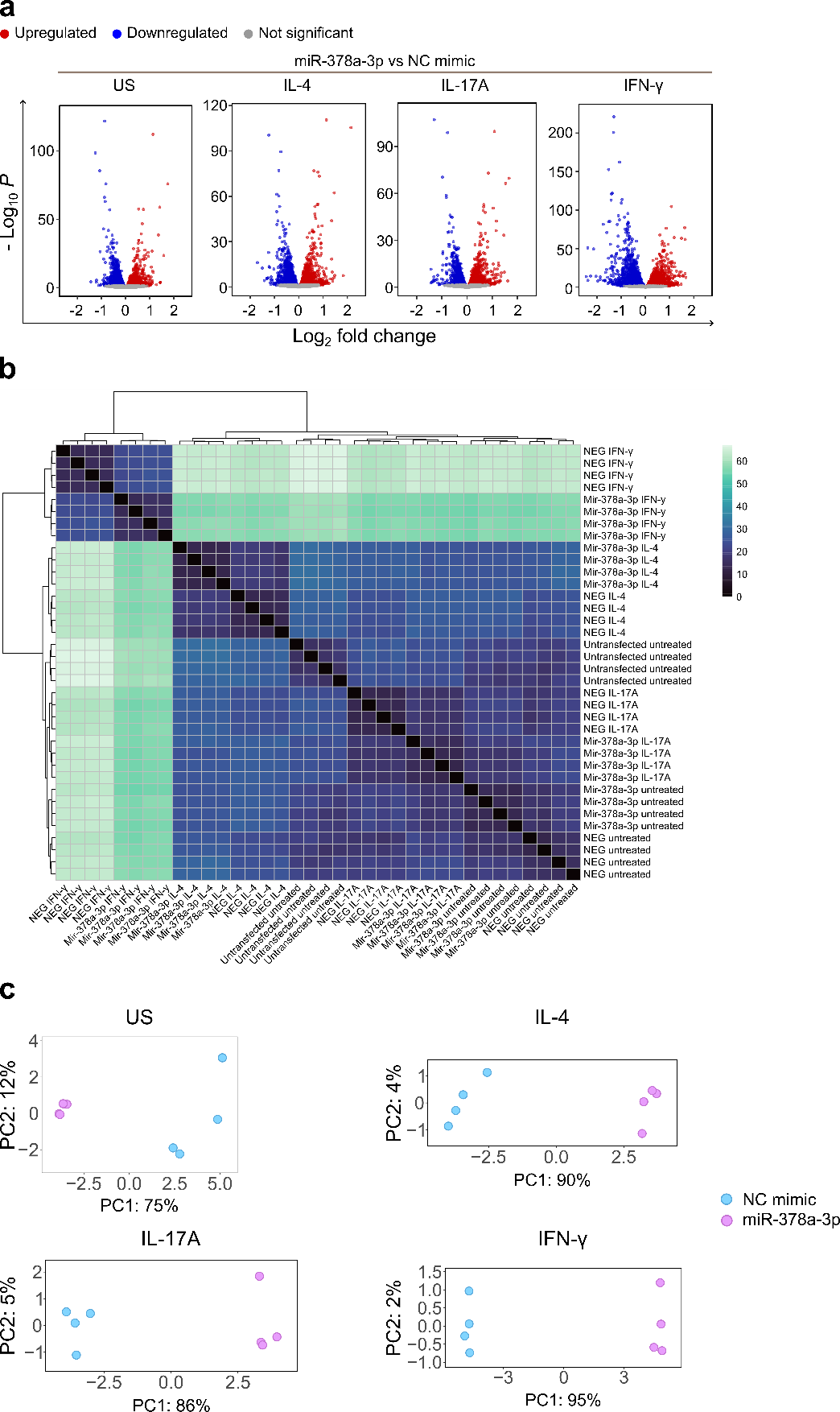

**Supplementary Figure S1.** miR-378a-3p overexpression and cytokine stimulation induce transcriptomic changes in primary keratinocytes. (a) Volcano plots of DEGs in NHEKs transfected with miR-378a-3p or negative control (NC) mimic under unstimulated (US) conditions or following stimulation with IL-4, IL-17A, or IFN-γ. DEGs were defined by baseMean ≥ 15 and FDR < 0.05. (b) Sample-to-sample distance heatmap of all RNA-seq samples across experimental conditions. NEG, negative control mimic. (c) Principal component analysis (PCA) of transcriptomic profiles of NC mimic and miR-378a-3p transfected NHEKs across the indicated experimental conditions.

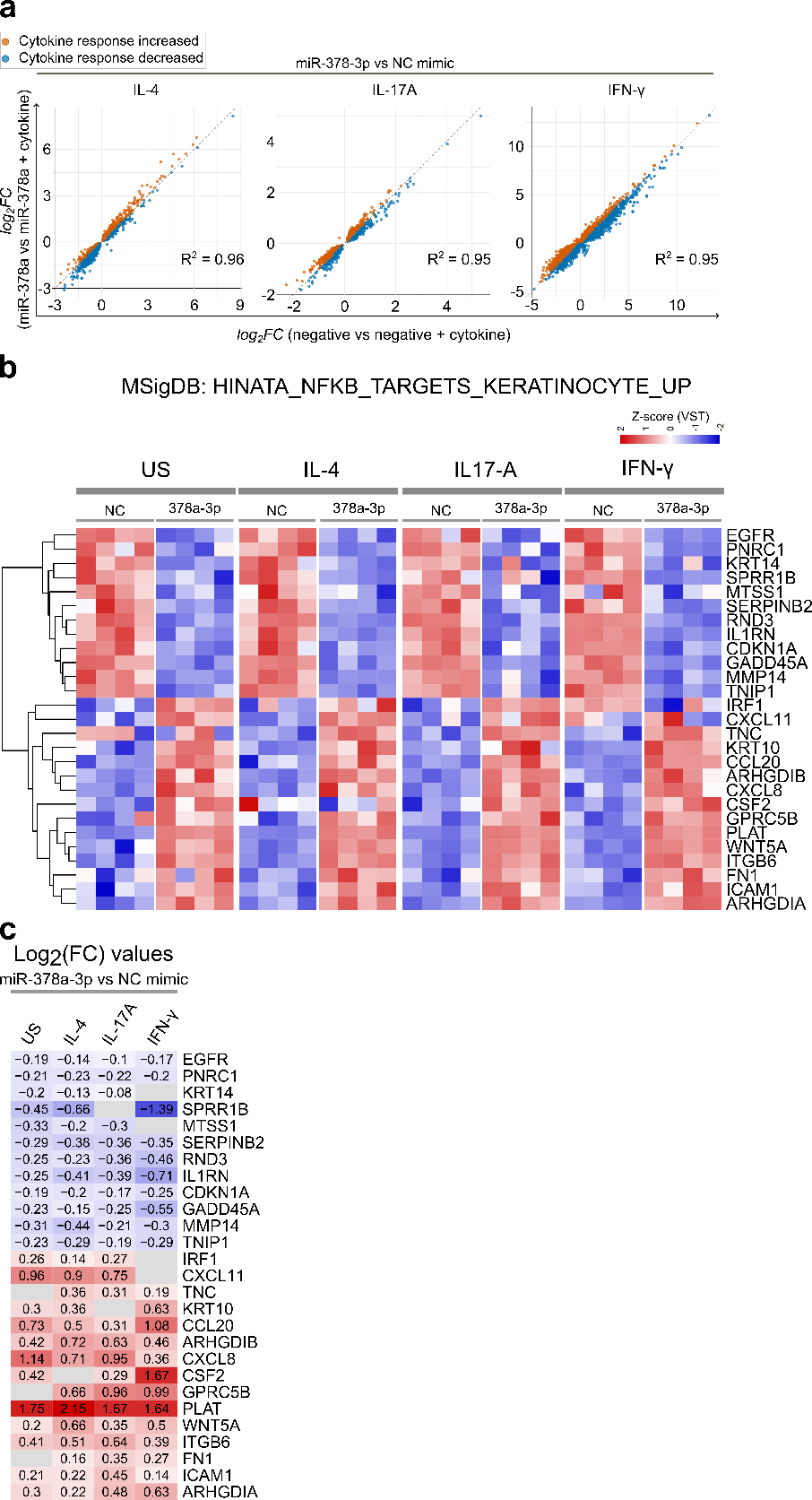

**Supplementary Figure S2.** miR-378a-3p overexpression influence on global cytokine responses and NF-κB target gene expression. (a) Scatter plots comparing cytokine-induced transcriptional landscapes between NHEKs transfected with miR-378a-3p and negative control (NC) mimic. For each condition, the log_2_ fold-change (stimulated versus unstimulated) in miR-378a-3p-transfected condition (y-axis) is plotted against the corresponding log_2_ fold-change in NC mimic-transfected condition (x-axis). Each dot represents one gene, the dashed line (y = x) indicates identical cytokine-induced responses between the two conditions. (b) Heatmap showing expression changes of keratinocyte-specific NF-κB target genes from the Molecular Signature Database (MSigDB: HINATA_NFKB_TARGET_KERATINOCYTE_UP) that were consistently modulated by miR-378a-3p overexpression in at least three conditions. (c) Log_2_ fold-change values for the genes shown in (b). Grey boxes indicate no change.

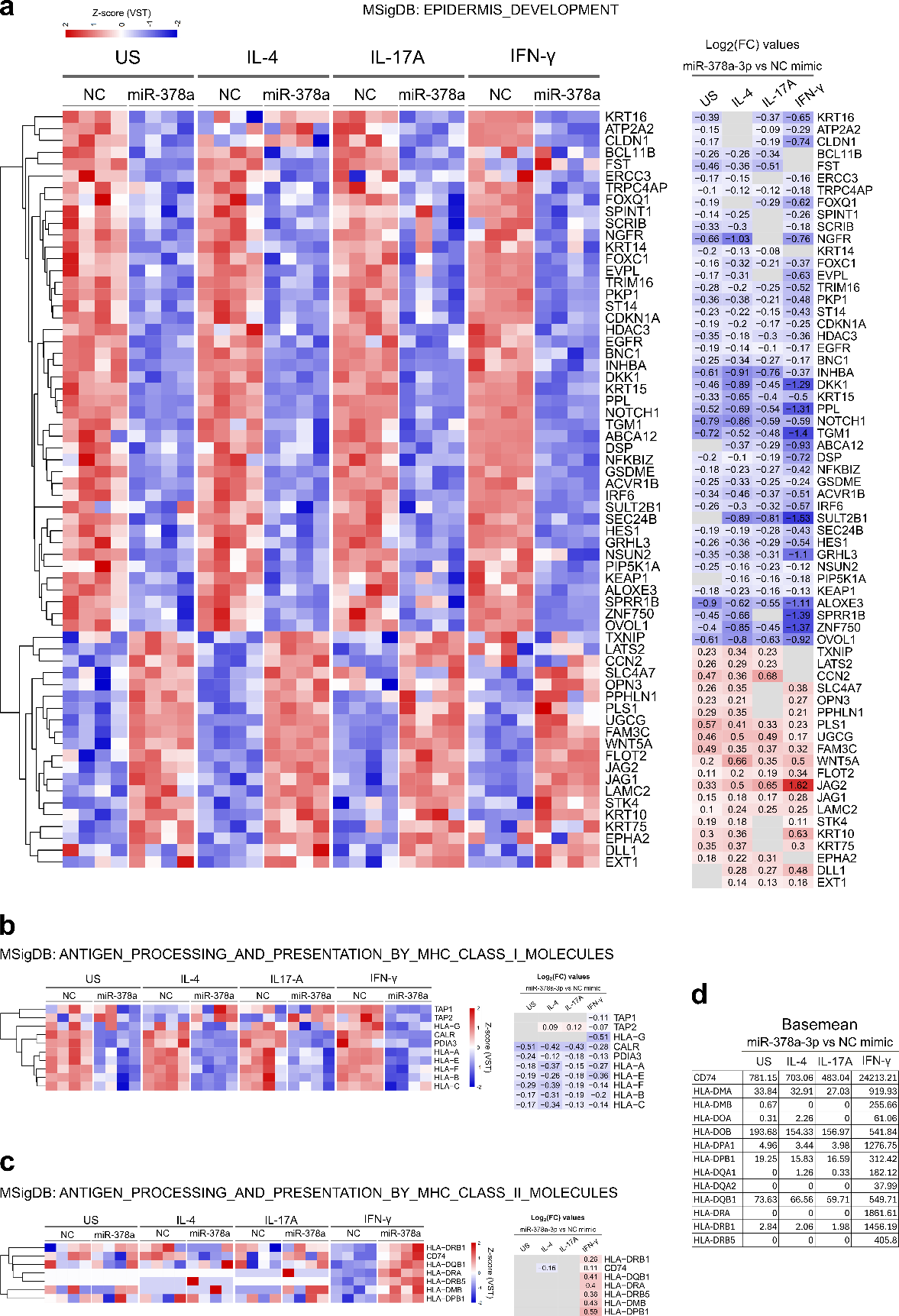

**Supplementary Figure S3.** miR-378a-3p modulates genes involved in epidermal development and antigen presentation. (a) Heatmap showing expression changes and corresponding log_2_ fold-change values for genes curated from the MSigDB (GOBP_EPIDERMIS_DEVELOPMENT) that were consistently modulated by miR-378a-3p across multiple conditions. (b, c) Heatmaps and log_2_ fold-change values for genes involved in major histocompatibility complex (MHC) class I (b) and MHC class II (c) antigen processing and presentation (MSigDB gene sets). Grey boxes indicate no change. (d) Expression levels (RNA-seq baseMean values) of MHC class II genes across all experimental conditions. This panel includes all genes from the curated MSigDB gene set, regardless of differential expression status.

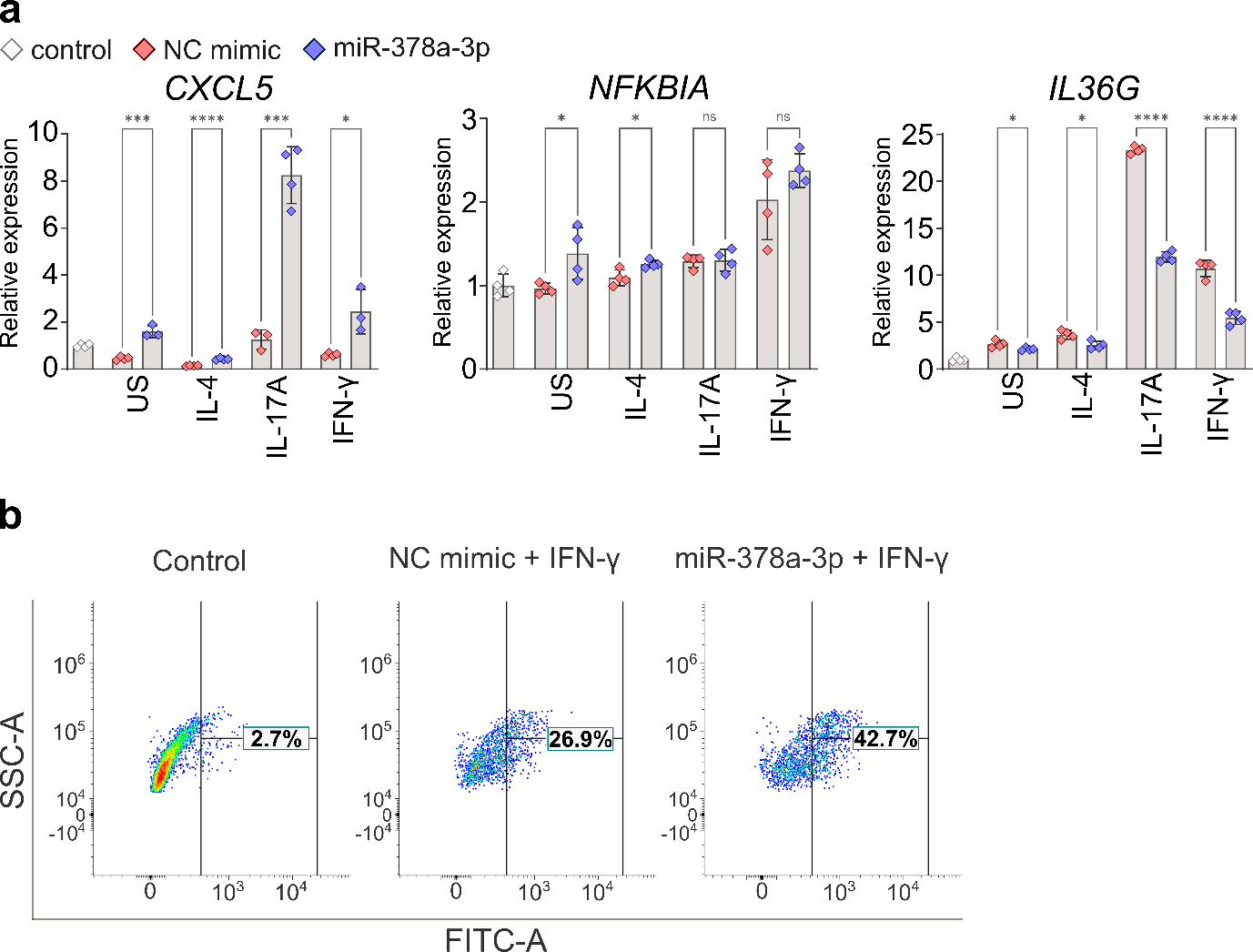

**Supplementary Figure S4.** miR-378a-3p overexpression alters the expression of mediators of inflammatory responses. NHEKs were transfected with miR-378a-3p or negative control (NC) mimic for 24 h and then stimulated with the indicated cytokines for an additional 24 h. Untransfected unstimulated NHEKs are designated as control. (a) RT-qPCR analysis of indicated genes. Data are presented as mean ± SD (n = 3-4). Unpaired two-tailed Student’s *t*-test. US, unstimulated; ns, not significant; **p* < 0.05, ****p* < 0.001, *****p* < 0.0001. (b) Flow cytometry analysis of HLA-DR (MHC class II) expression in indicated conditions showing the percentage of HLA-DR-positive cells.

**Supplementary Tables S3 and S4.**

**Supplementary Table S3.** Abbreviation for GSEA hallmark and Gene Ontology biological process terms used in Figure 2a and 2b.

- Inflammatory resp. – inflammatory response
- TNFα-NF-κB – TNFα signaling via NF-κB
- UPR – unfolded protein response
- EMT – epithelial-mesenchymal transition
- Estrogen resp. early/late – estrogen response early/late
- Resp. chemokine – response to chemokine
- Neutrophil chem. – neutrophil chemotaxis
- Humoral – humoral immune response
- GPCR – G protein-coupled receptor signaling pathway
- Epithelium dev. – epithelium development
- Organ morpho. – organ morphogenesis
- Skin epidermis dev. – skin epidermis development
- Granulocyte chem. – granulocyte chemotaxis
- Org. biosynth. pro. – organophosphate biosynthetic process
- Skin dev. – skin development
- Est. skin barrier – establishment of skin barrier
- Epidermis dev. – epidermis development
- Reg. gliogenesis – regulation of gliogenesis
- Granulocyte mig. – granulocyte migration
- Ubiquitin dep. pro. – regulation of ubiquitin-dependent protein catabolic process
- Fat cell diff. – fat cell differentiation
- Neg. re. cell dev. – negative regulation of cell development
- ECM regulation – regulation of extracellular matrix organization
- NMP biosynth. pro. – nucleoside monophosphate biosynthetic process
- rRNA metabolic pro. – rRNA metabolic process
- Ribosome biogen. – ribosome biogenesis
- Mito. gene expr. – mitochondrial gene expression
- Keratinocyte diff. – keratinocyte differentiation

**Supplementary Table S4.** RT-qPCR primer sequence used in this study.

| Gene | Forward (5’ – 3’) | Reverse (5’ – 3’) |
| --- | --- | --- |
| *CXCL8* | GCAGCTCTGTGTGAAGGTGCAGTT | TTCTGTGTTGGCGCAGTGTGGTC |
| *CCL20* | CGGCGAATCAGAAGCAAGCAA | GCATTGATGTCACAGCCTTCAT |
| *ICAM1* | TCAGTCAGTGTGACCGCAGA | GCGCCGGAAAGCTGTAGATG |
| *NFKBIA* | GATGTCAATGCTCAGGAGCCC | CCCCACACTTCAACAGGAGT |
| *CXCL5* | CAGACCACGCAAGGAGTTCA | TCTTCAGGGAGGCTACCACT |
| *IL1B* | AGAAGTACCTGAGCTCGCCA | TGGAAGGAGCACTTCATCTGT |
| *IL1RN* | CCCCATGGCTTTAGAGACGA | CCCAGATTCTGAAGGCTTGC |
| *IL18* | CTGCAGTCTACACAGCTTCGGG | TCCTGCGACAAATAGTTTGTTGC |
| *IL36G* | GGTGCTGAGACAACCACACT | ACAGCAACAGTGATTGATTGATAGA |
| *EEF1A* | CCACCTTTGGGTCGCTTTGCTGT | TGCCAGCTCCAGCAGCCTTCTT |
